# Probabilistic thresholding of functional connectomes: application to schizophrenia

**DOI:** 10.1101/233510

**Authors:** František Váša, Edward T. Bullmore, Ameera X. Patel

## Abstract

Functional connectomes are commonly analysed as sparse graphs, constructed by thresholding cross-correlations between regional neurophysiological signals. Thresholding generally retains the strongest edges (correlations), either by retaining edges surpassing a given absolute weight, or by constraining the edge density. The latter (more widely used) method risks inclusion of false positive edges at high edge densities and exclusion of true positive edges at low edge densities. Here we apply new wavelet-based methods, which enable construction of probabilistically-thresholded graphs controlled for type I error, to a dataset of resting-state fMRI scans of 56 patients with schizophrenia and 71 healthy controls. By thresholding connectomes to fixed edge-specific P value, we found that functional connectomes of patients with schizophrenia were more dysconnected than those of healthy controls, exhibiting a lower edge density and a higher number of (dis)connected components. Furthermore, many participants’ connectomes could not be built up to the fixed edge densities commonly studied in the literature (~5-30%), while controlling for type I error. Additionally, we showed that the topological randomisation previously reported in the schizophrenia literature is likely attributable to “non-significant” edges added when thresholding connectomes to fixed density based on correlation. Finally, by explicitly comparing connectomes thresholded by increasing P value and decreasing correlation, we showed that probabilistically thresholded connectomes show decreased randomness and increased consistency across participants. Our results have implications for future analysis of functional connectivity using graph theory, especially within datasets exhibiting heterogenous distributions of edge weights (correlations), between groups or across participants.

## 1. Introduction

Relationships between neurophysiological signals are thought to underlie communication in the brain (Fries, 2005); a complete description of such “functional connectivity” is called a functional connectome (Biswal et al., 2010). Functional connectomes are commonly analysed as sparse graphs, constructed by thresholding statistical associations (usually, correlations) between pairs of regional neurophysiological signals. Within graph theory, the relationships are called edges or links, while the brain regions are called nodes (Bullmore and Sporns, 2009). Thresholding retains the strongest edges, either by retaining edges surpassing a given absolute weight, or (more commonly) by ensuring that all connectomes have a fixed edge density, calculated as the ratio of the number of edges in the network to the total possible number of edges (Fornito et al., 2013). The resulting sparse graphs can subsequently be characterised using summary measures of topological organization, indicative of features such as network integration or segregation (Rubinov and Sporns, 2010).

As such measures are non-trivially dependent on the density of the underlying graph (van Wijk et al., 2010), weighted functional connectomes are traditionally thresholded to fixed edge density to enable comparisons of graph-theoretical measures across participants. Analyses using fixed-density thresholding are usually carried out across a range of thresholds to mitigate the fact that the choice of a single fixed density is arbitrary. While there have been recent efforts to determine a-priori thresholds (e.g. De Vico Fallani et al. (2017), based on the cost-efficiency trade-off (Bullmore and Sporns, 2012)), a statistically principled framework for thresholding individual graphs and analysing brain network connectivity has been lacking.

Recently, new wavelet-based methods have been proposed, which enable simultaneous denoising (Patel et al., 2014) and probabilistic inference (Patel and Bullmore, 2016) on functional connectomes constructed from individual subjects. These methods define the spatial variability in effective degrees of freedom (*df*) at each region or voxel after denoising and motion artefact removal, using wavelet despiking. These can then be used to convert edge statistics (correlations) into P values or probabilities (as described in Patel and Bullmore (2016) and Methods 2.2), thus enabling the construction of probabilistically-thresholded graphs. We note that application of P value thresholding to networks constructed using global denoising methods (such as scrubbing) and/or Pearson correlation unadjusted for effective *df* will not yield the same benefits as the methods applied here (see also Fig. 2 and Results 3.5), and will inflate type I error Patel and Bullmore (2016). Here we apply these new methods to resting-state fMRI data of patients with schizophrenia and healthy participants.

Schizophrenia has been described as a disorder involving both *disconnectivity* (Friston and Frith, 1995; Friston, 1998) and *dysconnectivity* (Bullmore et al., 1997; Stephan et al., 2006); the former referring to weaker or missing connections, the latter to aberrant connectivity more generally (Stephan et al., 2006; Fornito et al., 2012). Dysconnectivity in schizophrenia has been extensively studied using multiple methods, including graph theory (see e.g.: Bullmore and Sporns (2009) and Fornito et al. (2013) for reviews on analyses of neuroimaging data using graph theory, and Fornito et al. (2012) and van den Heuvel and Fornito (2014) for its specific applications to schizophrenia). These have generally reported global reductions in both structural (eg: van den Heuvel et al. (2010); Zalesky et al. (2011); Griffa et al. (2015)) and functional (Lynall et al., 2010; Zalesky et al., 2012; Lo et al., 2015) connectivity in schizophrenia. Some functional studies have shown localised increases in functional connectivity (Liu et al., 2008; Skudlarski et al., 2010), although these may reflect differences in preprocessing (e.g. use of partial correlations (Liu et al., 2008) or zero-centering of the correlation distributions (Skudlarski et al., 2010)). Furthermore, functional connectivity studies generally agree that brain networks are topologically altered in schizophrenia, although the nature of these changes and the specific topological measures used to assess them vary between studies (for review, see Fornito et al. (2012) and van den Heuvel and Fornito (2014)).

Among these studies, thresholding to fixed edge density has been more common. The popularity of fixed-density thresholding is likely due to the known dependence of “traditional” higher-order graph-theoretical measures on edge density (van Wijk et al., 2010). Several studies (Rubinov et al., 2009; Alexander-Bloch et al., 2010; Lynall et al., 2010) converge on topological alterations such as increased efficiency and/or reduced clustering, consistent with a subtle randomisation of the connectome in schizophrenia; this has been proposed as an endophenotype of the disorder (Lo et al., 2015). However, it has been hypothesized that this “randomisation” might result from the application of fixed-density thresholds to functional connectomes of patients with schizophrenia, presenting decreased edge weights: “in the presence of a global reduction of mean functional connectivity in patients […] any analysis of graphs matched for connection density, *κ*, will result in the inclusion of proportionally more low-value (non-significant) edges in patients’ networks. If these values merely reflect noise, their inclusion will produce a more random topology” (Fornito et al., 2012). A recent study demonstrated that group differences in mean functional connectivity (correlation) do indeed lead to group differences in topological organisation, and proposed to correct for between-group differences in mean functional correlation using regression or permutation (van den Heuvel et al., 2017).

Here, we present the first analysis of neuroimaging data using the methods for combined denoising and probabilistic inference presented in Patel and Bullmore (2016). We thresholded functional MRI scans of 56 patients with schizophrenia and 71 healthy controls (the COBRE dataset) based on statistical significance of edges, after taking into account the effects of motion and other artefacts. We aimed to investigate the general implications of probabilistic thresholding on graph theoretic analysis of brain network connectivity. In this context we used data from patients with schizophrenia to revisit the dysconnectivity and topological randomisation hypotheses using these new statistically principled thresholding methods. Specific questions included: (i) To what density can each participants’ connectome be reconstructed, while ensuring that all retained edges remain statistically significant after accounting for the effects of motion? Does this density have diagnostic potential? (ii) Conversely, what proportion of connectomes can be reconstructed up to fixed densities generally considered in the literature? (iii) (To what extent) are differences in topological organization sensitive to “non-significant” edges usually added when thresholding connectomes to fixed density?

## 2. Methods

### 2.1. MRI data and pre-processing

Raw anatomical and functional MRI scans of 72 patients with schizophrenia and 75 healthy controls were made available by the Mind Research Network and University of New Mexico (http://fcon_1000.projects.nitrc.org/indi/retro/cobre.html). Informed consent was obtained from all subjects according to institutional guidelines required by the Institutional Review Board at the University New Mexico (UNM).

All participants were scanned on a 3 Tesla SIEMENS TIM scanner. Structural data was collected using a multiecho MPRAGE (MEMPR) sequence with the following parameters: TR = 2.53 s, TE = [1.64, 3.5, 5.36, 7.22, 9.08] ms, TI = 900 ms, matrix size 256 × 256, 176 slices, voxel size =1×1×1 mm^3^. Resting-state data was collected with echo-planar imaging (EPI) TR = 2 s, TE = 29 ms, matrix size 64 × 64, 32 slices, voxel size = 3 × 3 × 4 mm^3^, scan duration = 304 s (152 volumes). Subjects were instructed to keep their eyes open during the scan. Subject ages ranged from 18 to 65 years old. Diagnostic information was collected using the Structured Clinical Interview used for DSM Disorders (SCID).

Core image processing of the structural and functional images was done in AFNI. This included: skullstrip of the structural image, slice timing correction, rigid-body head movement correction (to the first frame of data), obliquity transform of the functional to the structural image, affine co-registration of the functional to the structural image using a grey matter mask, non-linear standard space transform to the MNI152 template in standard space, spatial smoothing (6mm FWHM) and within-run intensity normalisation to a whole-brain median of 1000. Subsequent time series denoising steps included: voxel-wise wavelet despiking using the BrainWavelet Toolbox (www.brainwavelet.org), segmentation of CSF signal using FSL fast, and linear regression of 6 movement parameters, their first-order derivatives and CSF signal. All pre-processing steps were performed using AFNI software (Cox, 1996), except CSF segmentation which was performed using FSL FAST (Smith et al., 2004), and time series de-noising which was done using the BrainWavelet Toolbox for denoising motion artefacts (Patel et al., 2014).

Following data processing, participants with a mean spike percentage (the percentage of gray mater voxels containing a motion-related spike in the wavelet domain, averaged across time points) greater than 7.5% were excluded from further analysis. This threshold was chosen as it was the highest threshold that enabled the elimination of a between-group bias in motion and average effective *df* (after subject exclusion). This resulted in the exclusion of 15 patients with schizophrenia and 3 healthy controls. Additionally, one patient was excluded due to a truncated run during acquisition, and one healthy control due to reconstruction errors. Thus, 56 patients with schizophrenia and 71 healthy controls were included in the study. The framewise displacement (FD) in the two groups, expressed as median [first, third quartiles] ([Q1,Q3]) was 0.27 [0.20, 0. 40] in healthy controls, 0.35 [0.22, 0.47] in patients with schizophrenia.

### 2.2. Wavelet despiking and estimation of effective df

Wavelet despiking is a method for voxel-wise spatially-adaptive denoising of motion artefacts across frequencies that accounts for the highly non-linear nature of these artefacts. The first step of the algorithm is to perform a maximal overlap discrete wavelet transform (MODWT) on the time series (length = *t*) from each voxel which derives a set of frequency bands (or scales = *j*) from the signal creating a set of *t × j* wavelet coefficients. Denoising is conducted in the time-scale plane on these *t × j* coefficients for each voxel. The algorithm identifies chains of maximal and minimal wavelet coefficients across frequencies and splits the coefficients for each voxel time series into two additive sets: one representing noise coefficients (Φ), the other representing non-noise or ‘signal’ coefficients (*α*), as described in equation 1:

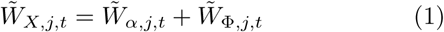

where 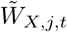 is the MODWT of voxel time-series *X_t_*.

In the final step, the algorithm recomposes the denoised time series from *α*, setting all Φ = 0 (a process known as hard thresholding), to yield a denoised time series of length *t*.

Effective degrees of freedom (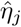 or simply *df*) are estimated for each scale in the time-scale plane from the ‘signal’ coefficients, *α*, using equation 2 below, as described in Patel and Bullmore (2016); *M* is the number of non-boundary coefficients (please see Patel and Bullmore (2016) for a full description).

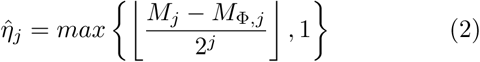

This results in estimation of 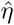 (*df*) for each voxel at each wavelet scale.

### 2.3. Graph construction

Individual networks were constructed using a parcellation of cortex into 470 nodes (Patel and Bullmore, 2016). 50 nodes were excluded due to incomplete coverage between subjects and dropout, defined as regions with insufficient signal coverage across the full cohort. Edge weights were calculated as Pearson correlations in the wavelet domain between the remaining 420 nodes, separately for each wavelet scale. Based on previous work indicating that wavelet scale 2 (0.060-0.125 Hz) is the most sensitive to differences between patients with schizophrenia and healthy controls (Lynall et al., 2010), the results primarily focus on this scale; still, the main analyses were repeated at scales 1 and 3 (see Supplementary Information). For a schematic representation of the graph construction pipeline, see Fig. 1A.

**Figure 1:**
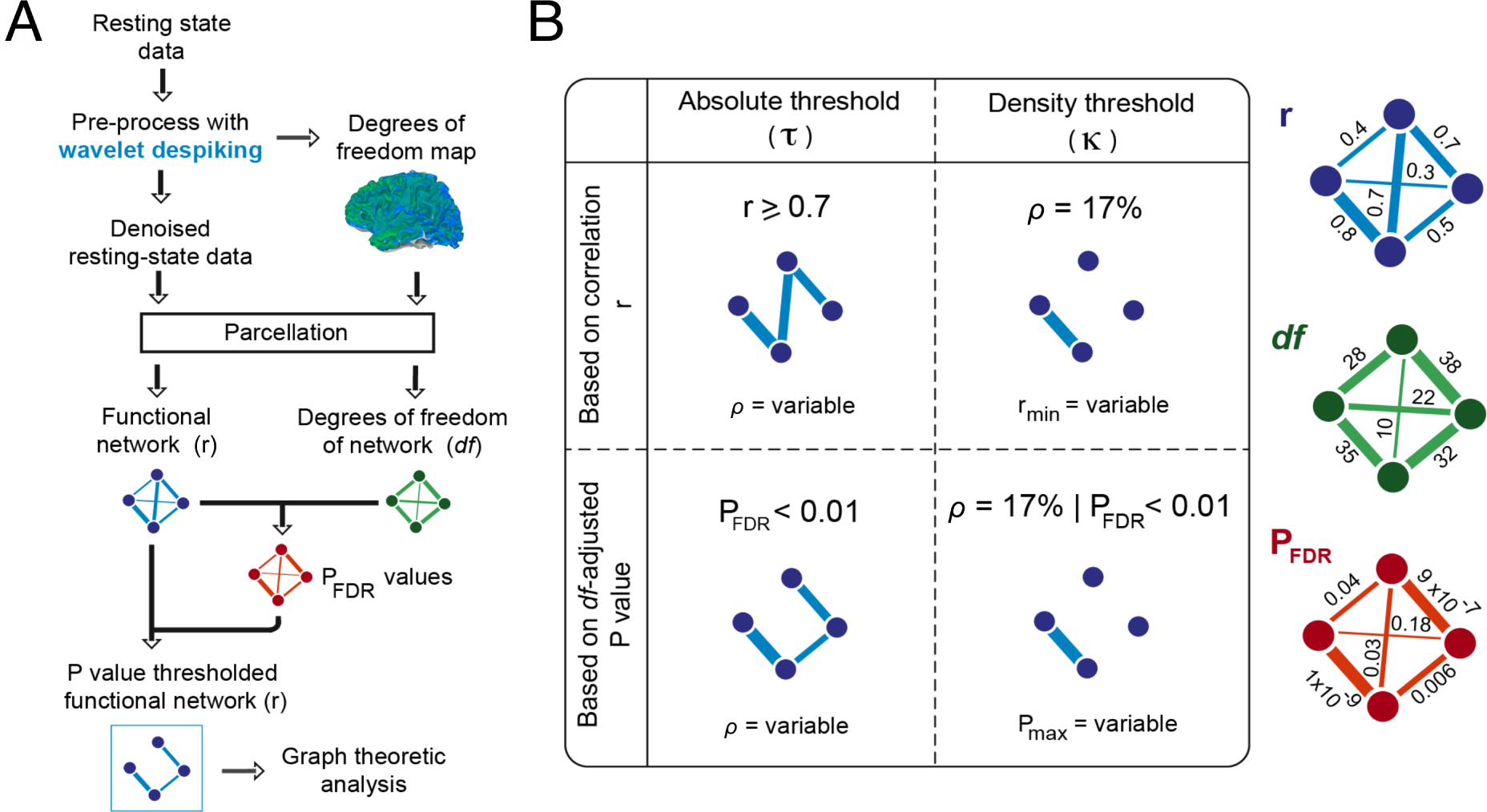
Methods of probabilistic graph construction and analysis. A) Pipeline for graph construction using wavelet *df*. During pre-processing and denoising with wavelet despiking (Patel et al., 2014), voxel-wise effective degrees of freedom (*df*) are extracted (Patel and Bullmore, 2016). Following parcellation of cortex into nodes and construction of a functional correlation network, edge-specific *df* values are used to obtain edge-specific P values, which (following FDR adjustment for multiple comparisons) can be used to threshold the network. B) Illustration of potential differences between graphs thresholded based on correlations, or *df*-adjusted P values. Graph thresholding, either to variable edge density using an absolute threshold (*τ*; first column) or to fixed edge density using a proportional threshold (*κ*; second column) has traditionally been performed using correlation (first row). However, application of analogous thresholding methods using *df*-adjusted P values (second row) can lead to different topologies.

Subject-specific *df* maps were parcellated using the same template used for parcellating the time series. This gave regional (or nodal) *df* estimates for each wavelet scale. When constructing graphs from this set of nodal *df,* edges were assigned the minimum *df* of the connecting nodes. So if node 1 had *df* = 30, and node 2 had *df* = 40, the edge *df* connecting these nodes was assigned *df* = 30. The correlation (r) values between nodal time series were then converted to 2-tailed P values using the Fisher r-to-Z transform and comparing to the standard normal distribution, normalising for edge *df* (equation 3).

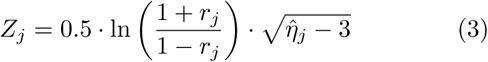

Unthresholded functional connectomes were characterised using the global mean correlation and node strength, calculated respectively as the mean of the upper triangular parts and rows (or equally, columns) of each participant’s adjacency matrix. Positive and negative correlations were not separately considered in this analysis.

### 2.4. Probabilistic thresholding methods

Throughout this paper, we use the terms “probabilistic thresholding” and “P value thresholding” (or “P-thresholding”) interchangeably, as the P value is an estimate of connection probability. (Specifically, it is the probability of correctly rejecting the null hypothesis, or of obtaining an observed edge test statistic at least as extreme as the one we observed, if the experiment were to be repeated.) We use the term “significant” to mean FDR-adjusted P < 0.01. However, we note that the method itself is deterministic, in that for a given network and threshold, the same edges will be retained.

An illustration of differences in topological organisation that may arise when thresholding connectomes based on correlations or P values is shown in Fig. 1B.

#### 2.4.1. Fixed P value thresholding

First, we thresholded each participant’s graph at an FDR-adjusted significance level of *α* = 0.01 - an approach which bears some similarity to (absolute) weight-based thresholding (Fornito et al., 2013) but additionally controls for type I error (“false positives”; Patel and Bullmore (2016)). We term all edges surviving this probabilistic threshold “significant edges”. Very few negatively-weighted edges survived (within and across participants), and inclusion of these edges had a negligible effect on downstream topological measures (see Supplementary Information). Negative edges were therefore excluded from the main analysis.

To study topological disconnectivity, P-thresholded connectomes for individual subjects were binarised by assigning all retained edges a uniform weight of 1. Upon thresholding and binarisation, many of the individual connectomes fractionated into multiple components. Therefore, we studied the topological disconnectivity within each participant’s connectome as a signal of interest, evaluating it using several global summary measures:

1. Edge density: the ratio of the number of edges present in the P-thresholded graph to the total number of possible edges.
2. Connected components analysis: these are sub-graphs within which any node is reachable from any other node by a path. We evaluated both the number of connected components, and the size of the largest component for each participant.
3. The percolation threshold: this is the threshold that determines connectedness of all nodes. In this case it is the (FDR-adjusted) P value below which all nodes form part of a single “giant” connected component, while above it the graph begins to fractionate into multiple (dis)connected components.

Additionally, we calculated the average Euclidean distances spanned by retained edges.

Further, we assessed local measures analogous to the global ones – node degree (the number of edges connected to a node), local edge density (degree normalised by number of edges in the graph), a nodal connected component score (defined as the size of the component the node is connected to, normalised relative to the largest component), and average distance spanned by a node’s edges.

Summary measures were compared between groups using two-tailed Mann-Whitney U (MWU) non-parametric tests. Effect sizes were quantified using the “simple difference formula” which results in a rank-biserial correlation r. All possible combinations of pairs of measurements from the two groups are classified as either favourable (*f*) or unfavourable (*u*) to the null hypothesis; then, the rank-biserial *r* is calculated as the difference between the two (*r* = *f* - *u*) (Kerby, 2014). The rank-biserial correlation is a (signed) value between 1 and -1; we used healthy controls as the reference population, resulting in positive values of *r* for measures that are increased in schizophrenia, and negative values of *r* for decreases in schizophrenia.

Finally, we examined whether functional connectomes thresholded using fixed P values exhibit some of the known hallmarks of topological organisation, including the presence of highly connected hub nodes (Sporns et al., 2007) and decomposability into densely intra-connected but sparsely inter-connected modules (Sporns and Betzel, 2016).

#### 2.4.2 Fixed density thresholding

In a second analysis, we thresholded individual connectomes to fixed edge density (from 1 to 35%, in steps of 1%), adding edges in order of increasing P value. In doing so, we studied the evolution of the maximum *df*-corrected P value per participant as a function of edge density. This analysis also indicates the maximum edge density that one can add edges to by P value for a given participant – once a threshold of P = 1 is reached, all remaining edges are equally unlikely and no further edges may be added by P value. Furthermore, we asked what proportion of our participants’ connectomes could be built up to a given fixed edge density, under the condition that all *df*-corrected P values remain statistically significant (P_FDR_ < 0.01). In adding edges in order of increasing P value beyond a threshold of P = 0.01, an increasing number of negative edges might be added (SI Fig. S2). However, negative edges were again excluded as they had no qualitative impact on the results.

For each individual’s fixed-density graph, we assessed topological integration and segregation. Global integration was quantified using the global efficiency, which is defined as the average of the inverse shortest path length (Latora and Marchiori, 2001). Global segregation was quantified using transitivity, which is the ratio between numbers of triangles and connected triples of nodes in the network (Newman, 2010). We note that the interpretation of transitivity as a measure of topological segregation relies on its application to sparse networks (here, 1-35% edge density). Transitivity is maximal in the extreme case of a fully connected graph, in which one might also say that integration is maximal. We did not enforce node-connectedness (e.g. with minimum spanning tree (MST) thresholding (Alexander-Bloch et al., 2010; Fornito et al., 2016)), as doing so would introduce non-significant edges. However, we quantified the number of connected components, for each participant and at each edge density.

Using these measures, we investigated whether topological randomisation in schizophrenia reported in the literature may be driven by inclusion of non-significant edges (Fornito et al., 2012), which we define as any edge with Pfdr ≥ 0.01. This analysis was conducted in 3 parts:

1. We first compared the topology between groups at fixed connection densities, adding edges to individual connectomes by increasing P value (until we reached P = 1), but disregarding whether we had included non-significant edges. This was our comparator to the literature where connectomes are commonly thresholded to fixed density regardless of the statistical significance of edges. We expected that the schizophrenia group would show greater randomisation.
2. We then repeated the analysis in (1), but at each density, we only included subjects where all edges satisfied P_FDR_ < 0.01 (i.e. where all edges were significant by our definition). We expected the topological randomisation effect to disappear, implying that it is driven by inclusion of non-significant edges.
3. Finally, at each density, we subdivided both the schizophrenia and control groups into two subgroups: those subjects with non-significant edges at that density, and those without. Here we aimed to show definitively that it is inclusion of non-significant edges that drives the randomisation effect.

Group differences were again evaluated using two-tailed Mann-Whitney U tests, and effect sizes using the simple difference formula (Kerby, 2014). For analyses (2) and (3) above, as group sizes varied as a function of density, we used permutation testing to ensure our results were not driven by differences in statistical power. For analysis (2), we sampled *N* subjects from each whole group (i.e. controls or schizophrenia patients with all edge P < 1 at a given density) without replacement, where N*N* is the number of subjects used for the analysis at that density, and evaluated the effect size. For analysis (3), we shuffled group labels. In both cases, the permutation was run 10,000 times, and permutation-test P values (P_perm_) were calculated as the proportion of effect sizes of randomly sampled (or permuted) participants that exceeded the empirical effect size.

Finally, we repeated the above fixed-density analyses using values of efficiency and transitivity which were normalised with respect to their average values in 100 randomised networks with preserved degree distributions (Rubinov and Sporns, 2010), within each participant and at each density. Additionally, due to the previously reported relationships between group differences in mean correlation and topological measures (van den Heuvel et al., 2017), we evaluated differences in mean (unthresholded) correlation between subsets of participants whose edges were or were not significant at each density.

### 2.5 Comparison of probabilistic and correlation-based thresholding

Our final analysis consisted in comparing probabilistic thresholding, where edges are added in order of increasing P value (adjusted for effective *df*), and correlation-based thresholding, where edges are added in order of decreasing correlation. We note that in time-series with fixed nominal degrees of freedom (equal to the number of time points, which is an overestimate of the true effective *df*), the two approaches will lead to identical results, due to the perfectly monotonic relationship between the correlation coefficient and the P value (Fig. 2A). However, following adjustment for effective *df,* the P values become *df*-dependent – and thus different edges may be added in order of increasing P and decreasing r (Fig. 2B).

**Figure 2:**
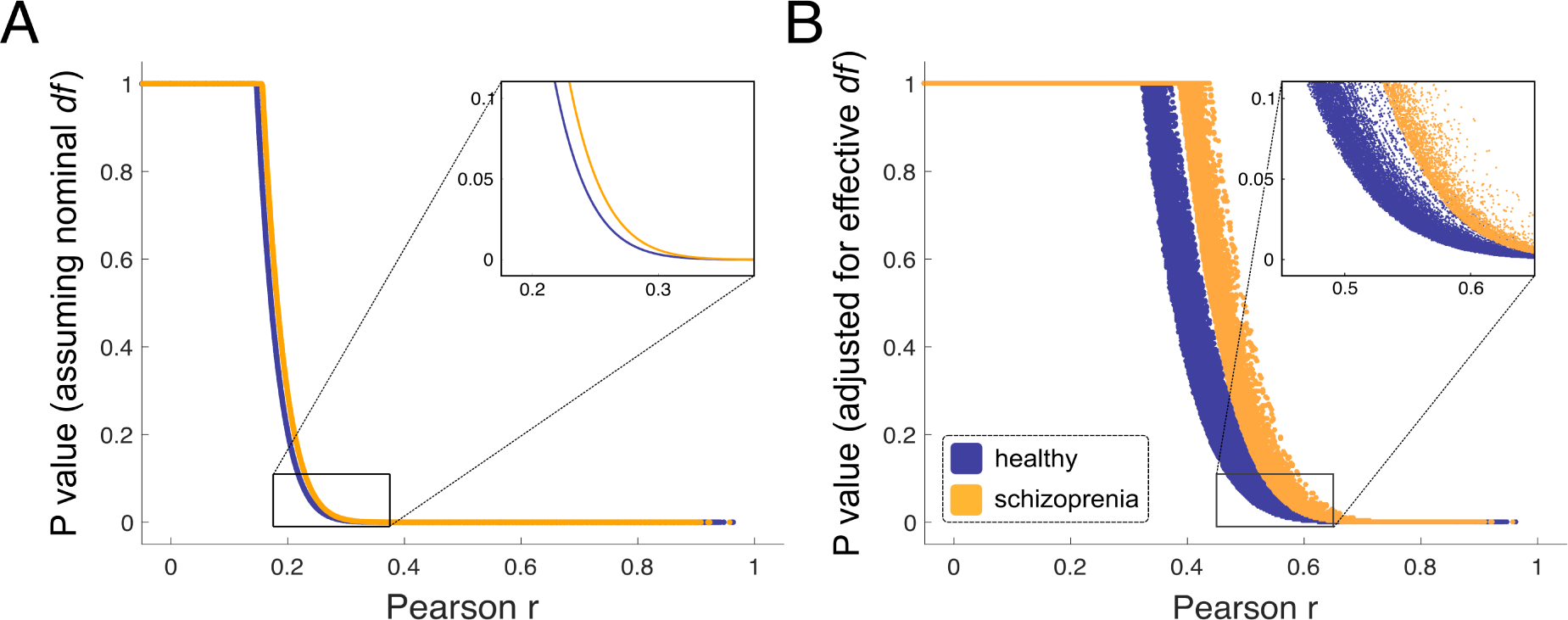
Impact of adjustment of P values for effective *df*. Illustration of the effect of adjustment of P values for effective *df*, on one example participant from each group. A) Without adjustment of P values for effective *df*, the relationship between the Pearson correlation and the P value is perfectly monotonic; thus, identical edges are added to the graph in order of decreasing r or increasing P. B) However, following adjustment of P values for effective *df*, different edges may be added in order of decreasing r and increasing P.

To compare the two approaches explicitly, we constructed for each participant a set of fixed-density graphs (in the range from 1 to 35% edge density) by adding edges in order of decreasing correlation coefficient r. We then compared these to networks constructed in order of increasing P value (whose construction is described above) using several approaches.

We first evaluated the proportion of edges that differ between the two networks, within participants and as a function of edge density.

Further, we evaluated differences in global efficiency and transitivity as a function of edge density. Within-group differences between topological measures derived from P or r-based thresholds were evaluated using the Wilcoxon signed-rank test (WSR; a non-parametric test of paired sample differences). We repeated these analyses using values of efficiency and transitivity which were normalised with respect to their average values in 100 randomised networks with preserved degree distributions (Rubinov and Sporns, 2010), for each thresholding method (increasing P or decreasing r), within each participant and at each density.

Finally, we examined whether thresholding by P value results in greater consistency of edges compared to thresholding by correlation. For each thresholding method (r and P) and at each edge density, we quantified the consistency of each edge edge by counting the number of times each particular edge was found across each group (controls and patients). We then converted this data to histograms by counting the number of edges that were observed for each level of consistency. Finally, we assessed the difference in these histograms of consistency between rand P-thresholded networks. The significance of any observed differences was determined by repeating the above procedure 10’000 times, following permutation or r- and P-thresholded networks within group; a P value was calculated as the proportion of permuted differences surpassing the empirical difference. The procedure is illustrated in Fig. 7A. As in previous analyses where networks were thresholded by fixed P value, we excluded participants whose connectomes contained edges with P = 1.

## 3. Results

### 3.1 Edge weight (correlation) distributions

The inter-regional wavelet correlation coefficients, or edge weights, were normally distributed and predominantly positive in both groups. Mean edge weights were significantly greater in the control group (mean weight: median = 0.37, first and third quartiles ([Q1,Q3]) = [0.29,0.45]) than in the schizophrenia group (mean weight: median = 0.27, [Q1,Q3] = [0.20,0.34]; rank-biserial *r* = -0.45, P_MWU_ = 1.2 · 10^−5^; Fig. 3A). The edge weight distribution for the schizophrenia group included more negative correlations than the control distribution.

**Figure 3:**
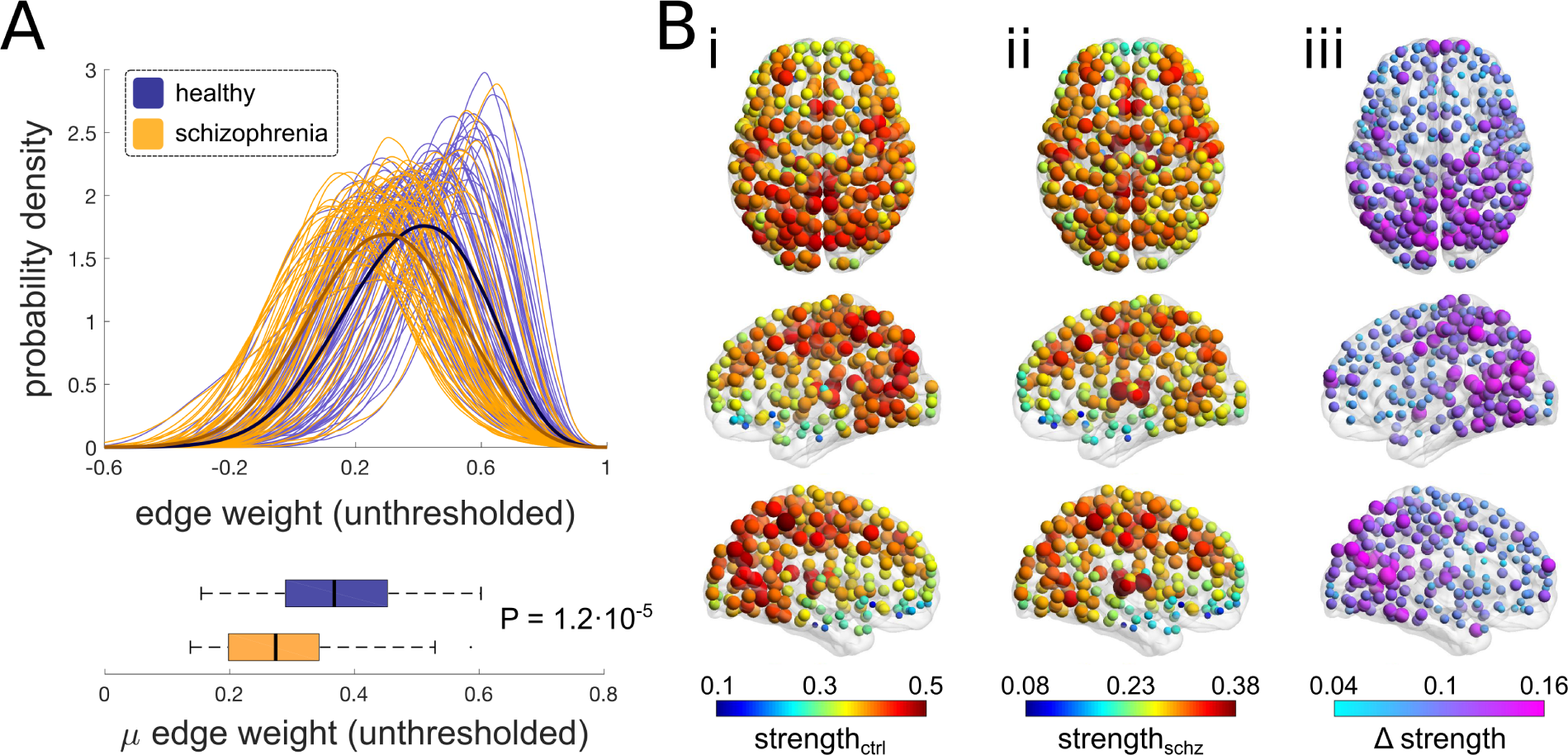
Decreased (unthresholded) correlations in patients with schizophrenia. A) Edge weights in unthresholded networks, as distributions within individual participants (thin lines) as well as averaged within groups (bold lines; top), and averaged within participants (bottom). Distributions were constructed using kernel density estimates. B) Maps of nodal connectivity strength, calculated as average correlation per node. (i) Median node strength of healthy controls, (ii) median node strength of patients with schizophrenia and (iii) difference in median node strength between groups (controls – patients). In map (iii), only the 389 nodes showing a significant difference are shown (Xia et al., 2013)

At a nodal level of analysis, edge weights were significantly reduced in the schizophrenia group compared to the control group at 389 nodes (FDR-adjusted P_MWU_ < 0.01). These nodal differences were attributable to greater nodal connectivity strength in healthy controls compared to patients with schizophrenia. There were no significant differences in nodal connectivity strength attributable to greater connectivity strength in participants with schizophrenia compared to healthy controls.

### 3.2 Properties of probabilistically-thresholded connectomes

We found that functional connectomes of patients with schizophrenia (thresholded at P_FDR_ < 0.01) were significantly more disconnected than those of healthy controls, exhibiting a lower edge density (rank-biserial *r* = -0.47, P_MWU_ = 6.5 · 10^−6^; Fig. 4Ai), higher numbers of connected components (rank-biserial *r* = 0.38, P_MWU_ = 2.1 · 10^−4^; Fig. 4Aii), and consequently a higher percolation threshold (rank-biserial *r* = 0.20, P_MWU_ = 0.052). In the schizophrenia group, the median connection density of these probabilistically-thresholded graphs was 4.0%, [Q1,Q3] = [2.6%,7.9%]; the median number of connected components was 17, [Q1,Q3] = [9,30]; and the median percolation threshold P value was 0.57, [Q1,Q3] = [0.16,1]. In the equivalently thresholded graphs for the control group, the median connection density was 13.8%, [Q1,Q3] = [4.7%,28.4%]; the median number of connected components was 8, [Q1,Q3] = [3.25,17.75]; and the median percolation threshold P value was 0.27, [Q1,Q3] = [0.069,0.82].

**Figure 4:**
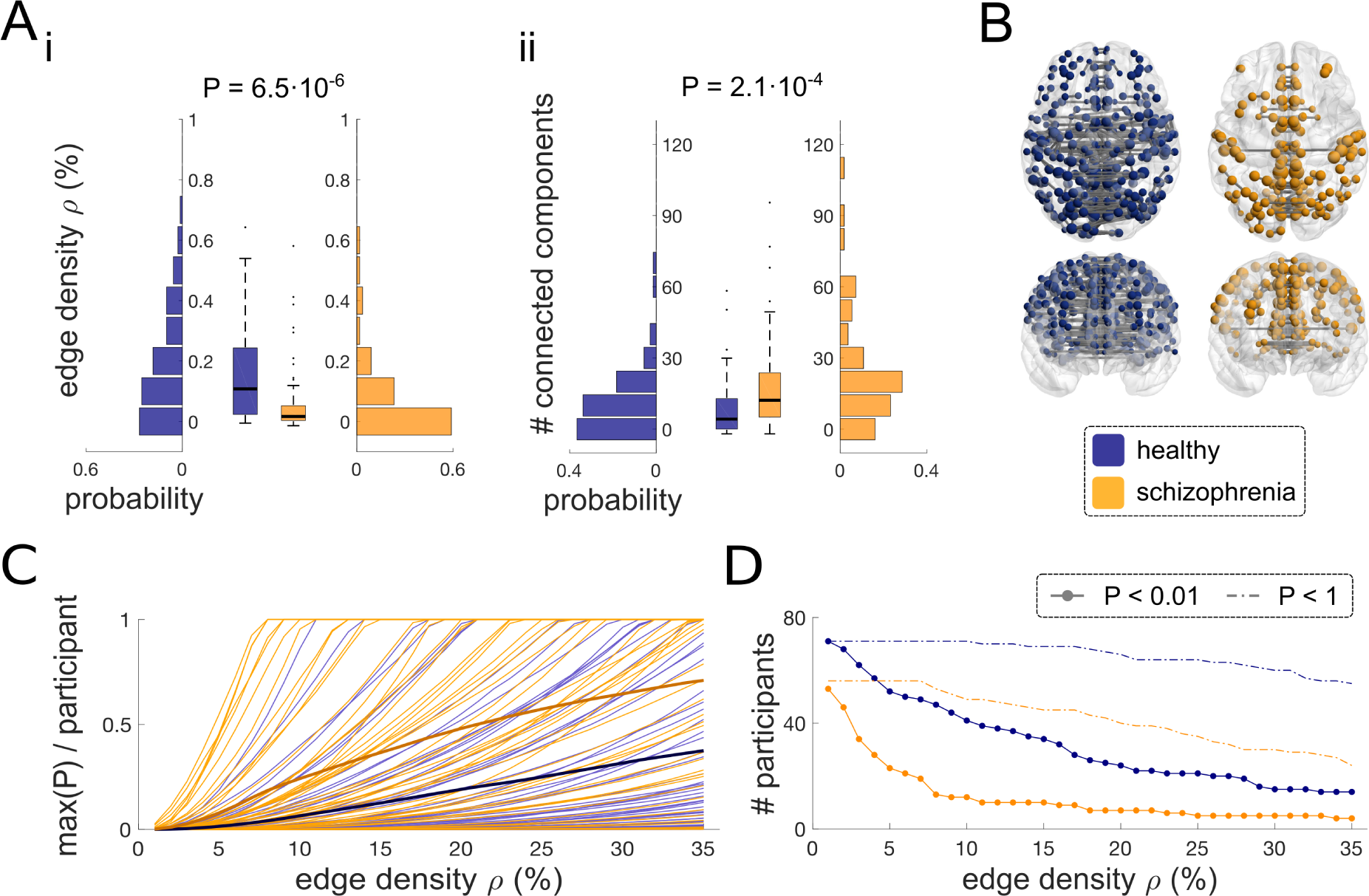
Connectomes thresholded based on fixed P value are more disconnected in patients with schizophrenia. A) Differences in i) edge density and ii) the number of connected components, with P values corresponding to two-tailed Mann-Whitney U tests. The x-axes indicate histogram frequency normalised by the number of participants (in each group). B) Illustrations of reduced edge density in schizophrenia, depicted as edges present in at least 90% participants per group, using brain network plots (only nodes connected to an edge are visualized; Xia et al. (2013)). C) Maximal FDR-adjusted P values as a function of fixed edge density threshold, for individual participants (thin lines) and averaged within groups (bold lines). D) Number of participants whose edges satisfy P_FDR_ < 0.01 (continuous lines with markers) and P < 1 (dotted lines), as a function of edge density. Once a P value of P = 1 is reached, further edges cannot be added by P value as all remaining edges are equally unlikely.

The reduced edge density of connectomes of patients with schizophrenia is seen in a visualisation of significant edges present in at least 90% of participants in each group (Fig. 4B).

Edges in the probabilistically-thresholded networks spanned shorter distances in patients with schizophrenia (rank-biserial *r* = -0.45, P_MWU_ = 1.3 · 10^−5^). The median average connection distance in patients was 48.7 mm, [Q1,Q3] = [40.5 mm, 58.0 mm], while the median average connection distance in controls was 60.6 mm, [Q1,Q3] = [52.6 mm, 69.2 mm]).

Finally, the variable-density networks presented the same hallmarks of topological organisation as fixed-density networks. Both groups exhibited spatially heterogeneous degree distributions, with highly connected “hub” nodes located in parietal and occipital cortices (consistent with other studies that did not apply global signal regression, which tends to shift hubs from primary to association areas; Yan et al. (2013)). Similarly to (unthresholded) node strength, patients with schizophrenia exhibited decreased node degree (FDR-adjusted P_MWU_ < 0.01) at 382/420 nodes (SI Fig. S1A). However, these decreases were driven by group differences in edge density – when local connectivity was assessed using a measure of local edge density (degree normalised by the participants’ edge density) no nodes showed significant differences between the two groups. Similarly, the two groups showed no significant differences in the size of the connected component per node (normalised by the largest component in the graph). Furthermore, the P-thresholded graphs were decomposable into densely intra-connected but sparsely inter-connected modules using the Louvain community algorithm (Blondel et al., 2008). For a resolution parameter of *γ* = 1, a consensus modular organisation (Lancichinetti and Fortunato, 2012) across participants (100 runs per participant) within each group yielded four modules per group (SI Fig. S1B), which were highly similar – only 26/420 nodes were assigned to different modules in the two groups.

### 3.3 To what density can functional connectomes be built?

Next, we used probabilistic thresholding to address an important question: to what density can we build functional connectomes to avoid adding false-positive edges? Here, we built networks by adding edges in order of increasing P_FDR_ and thresholded to a fixed edge density (see Methods). We found that the largest edge P_FDR_ in any given subject’s network rose rapidly as the network density was increased from 1 to 35% (Fig. 4C). In other words, many of the graphs needed to be constrained at very low connection densities to prevent inclusion of non-significant edges; this was true for both groups. Accordingly, if we require all edges in any given subject’s network to have P_FDR_ < 0.01 (which we term “significant edges”), the number of subjects meeting this criterion drops off rapidly as edge density is increased. The drop-off is faster for patients with schizophrenia (Fig. 4D) meaning that the maximum connection density that graphs can be built to, while ensuring that all participants’ connectomes contain only significant edges (edges with P_FDR_ < 0.01), is much lower in the schizophrenia group. For example, the maximum edge density while ensuring that 95% of connectomes contain only significant edges is only 2% in the control group and 1% in the schizophrenia group. Similarly, the maximum edge density while ensuring that 50% of connectomes contain only significant edges is 13% in the control group and 4% in the schizophrenia group.

### 3.4. What can probabilistic thresholding tell us about measures of integration and segregation?

Next, we used probabilistic thresholding to address the hypothesis of topological randomisation in schizophrenia, by studying group differences in topological integration (global efficiency) and segregation (transitivity). As described in section 2.4.2, we first thresholded all participants’ networks to fixed edge density, disregarding whether this included non-significant edges or not (but only considering participant at densities at which their connectomes contained edges with P < 1). Consistent with the literature, patients with schizophrenia showed significantly increased efficiency, indicative of greater topological randomisation, across a range of densities (P_MWU_ < 0.05 at 2-35% density; Fig. 5A).

In the next analysis, at each density, we included only subjects where all edges met the criterion P_FDR_ < 0.01. In other words, we excluded subjects (at each density) whose connectomes contained non-significant edges. As expected, the difference in global efficiency between groups disappeared at nearly all densities (P_MWU_ < 0.05 at 5 and 7% density; Fig. 5B).

To identify whether it was indeed inclusion of nonsignificant edges which led to the spurious result of greater topological randomisation in schizophrenia, we subdivided each of our two groups into two further subgroups at each connection density: those that did not contain any nonsignificant edges (i.e. all edge P_FDR_ < 0.01), and those that did. The ratios of the size of these subgroups (as well as a third subgroup of participants whose remaining edges all display P = 1 and who were not considered in the analyses) are depicted in Fig. 5C and D (top row). We then compared the “significant” edge subgroup with the “non-significant” (or noisy) subgroup. We found that global efficiency was significantly increased in the group that contained non-significant edges compared to the group where all edges showed P_FDR_ < 0.01, across connection densities, for both healthy controls (both P_MWU_ and P_perm_ < 0.05 at 3-35% density; Fig. 5C) and patients with schizophrenia (P_MWU_ < 0.05 at 1-35% density, P_perm_ < 0.05 at 1% and 3-35% density; Fig. 5D). This suggests that it is in fact the inclusion of non-significant edges that artefactually inflates efficiency, through increased randomisation.

These results were generally consistent when normalised with respect to random networks (where values of randomised efficiency tend to 1, as expected; SI Fig. S3). Although differences in efficiency between full groups (participants with all edge P < 1) were reduced relative to un-normalised efficiency (PMWU < 0.05 at 22% and 33% edge density; SI Fig. S3A), subsets of non-significant participants showed substantial decreases in normalised efficiency relative to subsets of significant participants, both within healthy controls (P_MWU_ < 0.05 at 3-35% density, P_perm_ < 0.05 at 5-25% and 28-35% edge density; SI Fig. S3C) and within patients with schizophrenia (P_MWU_ < 0.05 at 1-35% density, P_perm_ < 0.05 at 1%, 3%, 13-15% and 16-35% edge density; SI Fig. S3D).

Patients with schizophrenia showed decreased transitivity at higher edge densities (P_MWU_ < 0.05 at 20-23% and 32-35% density) when comparing all participants with P < 1 (SI Fig. S4A). These differences disappeared when comparing subsets of significant participants (all P_MWU_ and P_perm_ > 0.05; SI Fig. S4B). Subsequently, transitivity was lower in the non-significant healthy controls (P_MWU_ and P_perm_ < 0.05 at 7-25% and 28-35% density; SI Fig. S4C), and in non-significant patients with schizophrenia (P_MWU_ and P_perm_ < 0.05 at 1% and 12-35% density; SI Fig. S4D). Conversely, when normalised with respect to random networks, patients with schizophrenia exhibited higher transitivity than healthy controls when comparing all participants with P < 1 (P_MWU_ < 0.05 at 1-34% density; SI Fig. S5A). When comparing subsets of significant participants, these differences were reduced (P_MWU_ < 0.05 at 1- 6% density; SI Fig. S5B). Furthermore, “non-significant” participants displayed higher transitivity than “significant” ones, both within healthy controls (P_MWU_ < 0.05 at 7-25% and 28-35%, P_perm_ < 0.05 at 2-35% density; SI Fig. S5C) and patients with schizophrenia (P_MWU_ < 0.05 at 1% and 12-35% density, P_perm_ < 0.05 at 1-35% density; SI Fig. S5D).

Furthermore, when all participants (with all edge P<1) were compared, patients with schizophrenia showed a lower number of (larger) connected components, across densities (P_MWU_ < 0.05 at 1-30% and 32% density; SI Fig. S6A). However, when subsets of significant participants were compared, these differences were less extensive (P_MWU_ < 0.05 at 1-10% density; SI Fig. S6B). Accordingly, non-significant participants displayed a lower number of (smaller) connected components than significant participants, both within healthy controls (both P_MWU_ and P_perm_ < 0.05 at 3-35% density; SI Fig. S6C) and within patients with schizophrenia (both P_MWU_ and P_perm_ < 0.05 at 1-35% density; SI Fig. S6D).

Finally, we observed differences in mean correlation (of unthresholded networks) between healthy controls and patients with schizophrenia (comparing only connectomes with all edge P < 1 at a given density) revealed differences across edge densities (P_MWU_ < 0.05 at 1-27%, 29-30% and 33-35% density; SI Fig. S7A). These effects largely disappeared when only subsets of significant participants were compared (P_MWU_ < 0.05 at 1-5% density; SI Fig. S7B). Further, non-significant participants displayed lower mean correlation, both within healthy controls (both P_MWU_ and P_perm_ < 0.05 at 2-35% density; SI Fig. S7C) and patients with schizophrenia (both P_MWU_ and P_perm_ < 0.05 at 1-35% density; SI Fig. S7D).

### 3.5. Comparing r and P based thresholding

When constructing correlations using an equal number of data points (i.e.: time points in fMRI time-series), and assuming that N(*df*) = N(time-points), the relationship between the Pearson correlation coefficient r and the corresponding P value is perfectly monotonic (Fig. 2A). In this case, adding edges to a graph in order of (i) decreasing r or (ii) increasing P leads to identical topologies. However, wavelet-despiked time series exhibit regional heterogeneity in effective *df* after denoising for motion, which means that the relationship between r and P isn’t perfectly monotonic anymore (Fig. 2B). In this case, different edges may be added in order of (i) decreasing r or (ii) increasing P.

#### 3.5.1 Differences in topology between r- and P-thresholded connectomes

We quantified the proportion of edges that differ between r- and P-thresholded connectomes, as a function of edge density (Fig. 6A). The difference lay between 0 and 10% of edges present at a given edge density, with an average of ~2.8% edges differing between r and P-thresholded connectomes at low edge densities, slowly decreasing to an average of ~1.2% edges differing at higher edge densities. As with previous analyses involving thresholding of connectomes to fixed edge density by increasing P value, we only included participants with all edge P < 1 at each density.

**Figure 5:**
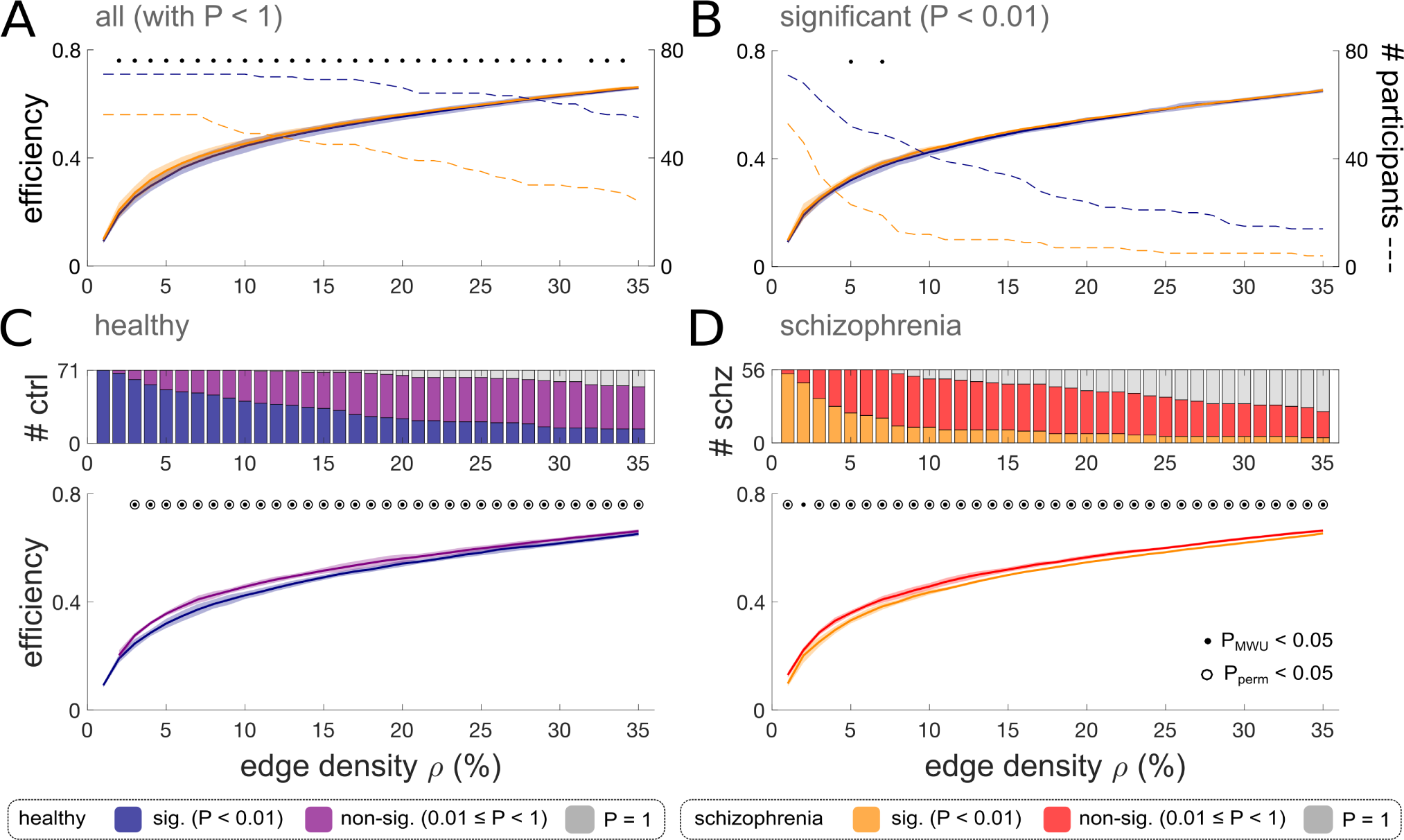
Effects on global efficiency of adding non-significant edges to functional connectomes. A) Global efficiency as a function of edge density, in all participants with edge P < 1 at each density (regardless of edge significance). B) Global efficiency as a function of edge density in the subset of participants whose edges are all significant at each density (left axis). The number of participants compared decreases as a function of edge density (right axis). C,D) Top: Density-dependent proportion of significant and non-significant participants, for healthy controls (C), and patients with schizophrenia (D). Bottom: Global efficiency as a function of edge density, between significant and non-significant healthy controls (C) and patients with schizophrenia (D). Black dots indicate P_MWU_ < 0.05, black circles indicate permutation test P_perm_ < 0.05. Thick lines indicate medians; shaded lines indicate quartiles. Effect sizes and P values of all two-tailed Mann-Whitney U tests are reported in SI Table S2.

**Figure 6:**
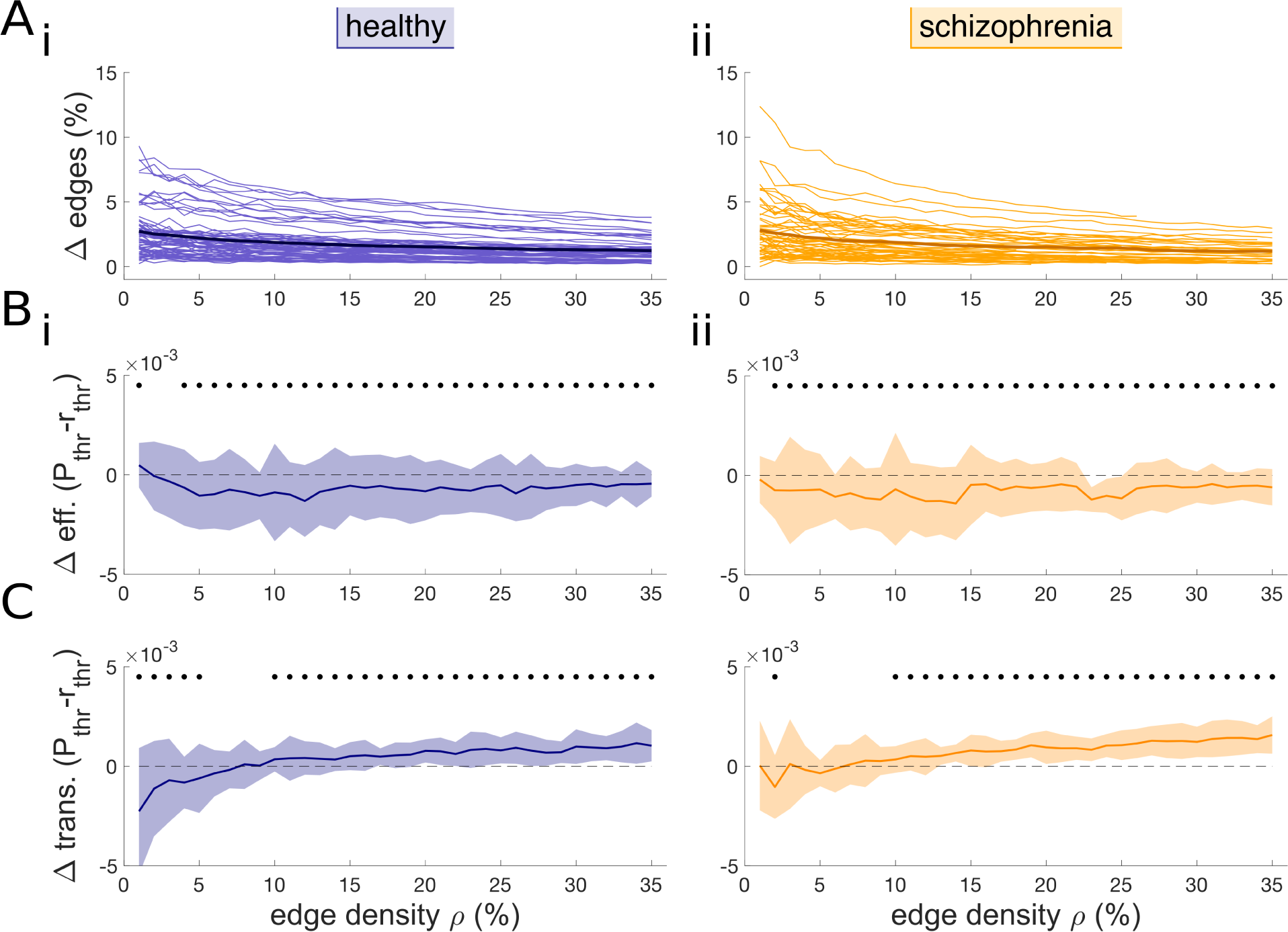
Differences in topology between networks thresholded by decreasing correlation or increasing P value. A) The percentage of edges that differs between functional connectomes thresholded by increasing P or decreasing r, as a function of fixed edge density, within (i) healthy controls and (ii) patients with schizophrenia. At low edge densities, ~2.8% edges differ between the two approaches, while at higher edge densities, this decreases to ~1.2% differing edges. B) Within-participant differences in efficiency between connectomes thresholded by increasing P and decreasing r. Plots represent the median difference in efficiency across participants. Efficiency is generally increased in connectomes thresholded by r (compared to those thresholded by P), in both healthy controls (i) and patients with schizophrenia (ii). C) Within-participant differences in transitivity between connectomes thresholded by increasing P and decreasing r. Plots represent the median difference in transitivity across participants. Transitivity is increased in connectomes thresholded by r (compared to those thresholded by P) at low edge densities, and increased at higher edge densities, in both healthy controls (i) and patients with schizophrenia (ii). Effect sizes and P values for results displayed in panels B and C are contained in SI Table S10.

Furthermore, we evaluated differences in measures of topological organisation between connectomes thresholded using increasing P and decreasing r. Global efficiency was weakly increased in r-thresholded networks relative to P-thresholded networks at a range of edge densities in both healthy controls (P_WSR_ < 0.05 at 4-35% density, Fig. 6Bi), and patients with schizophrenia (P_WSR_ < 0.05 at 2-35% density, Fig. 6Bii). Transitivity was lower in P-thresholded networks at low edge densities in both healthy controls (P_WSR_ < 0.05 at 1-5% density, Fig. 6Ci) and patients with schizophrenia (P_WSR_ < 0.05 at 2% density, Fig. 6Cii). However, at higher edge densities, transitivity was increased in P-thresholded networks relative to correlation-thresholded networks, in both healthy controls (P_WSR_ < 0.05 at 10-35% density) and in patients with schizophrenia (P_WSR_ < 0.05 at 10-35% density).

Results were highly consistent following normalisation of topological measures in r- and P-thresholded connectomes by corresponding average values from sets of 100 randomised networks with preserved degree distributions (Si Fig. S8).

##### 3.5.2 Consistency of edges in r- vs. P-thresholded connectomes

We further compared the consistency with which edges were present within both groups when thresholding in order of decreasing correlation or increasing P value (Fig. 7A). As in previous analyses where networks were thresholded by fixed P value, we excluded participants whose connectomes contained edges with P = 1. Exclusion of these subjects is indicated by a drop-off in the maximum possible consistency at higher densities (grey panels in top-right area of Fig. 7B).

We found a generally increased edge consistency in P-thresholded connectomes relative to r-thresholded ones, in both groups. The general pattern is of fewer edges being present in lower numbers of participants (blue bands on the left hand-side of Fig. 7B i and ii), and of more edges being present in a greater number of participants (a prevalence of light red on the right hand side of Fig. 7B i and ii); this is particularly evident at higher connection densities. However, while these differences were generally not significant (except at the lowest edge densities), there was an overall increase of consistency in P-thresholded networks compared to r-thresholded networks, as evidences by an increased frequency of edges absent from all participants when thresholding by P relative to r (left inset in Fig. 7B i and ii). This difference was significant at numerous edge densities, both within healthy controls (P_perm_ < 0.01 at 11-12%, 14-24%, 26-28%, 30% and 35%; Fig. 7Bi) and within patients with schizophrenia (P_perm_ < 0.01 at 9% and 13-35%; Fig. 7Bii).

#### 3.6 Sensitivity analyses

To rule out effects of age on our results, we evaluated within-group correlations between age and the average edge weight (of unthresholded connectomes) and the edge density and number of connected components (of connectomes thresholded by fixed P value). Furthermore, we investigated differences in these measures between male and female participants. We found no substantial evidence that these covariates affect our results. All correlations and P values are reported in supplementary table S1.

Furthermore, we investigated consistency of results across wavelet scales 1 (0.125-0.25 Hz) and 3 (0.03-0.06 Hz), as well across more lenient and stringent FDR-adjusted P value thresholds of *α* = 0.05 and *α* = 0.001. All results were qualitatively consistent with results reported above. For results of these analyses (including median values and interquartile ranges per group, as well as effect sizes and P values for group differences), see supplementary tables S2 and S3. At wavelet scale 4 (0.015-0.03 Hz), there were insufficient degrees of freedom for the addition of even a single edge.

**Figure 7:**
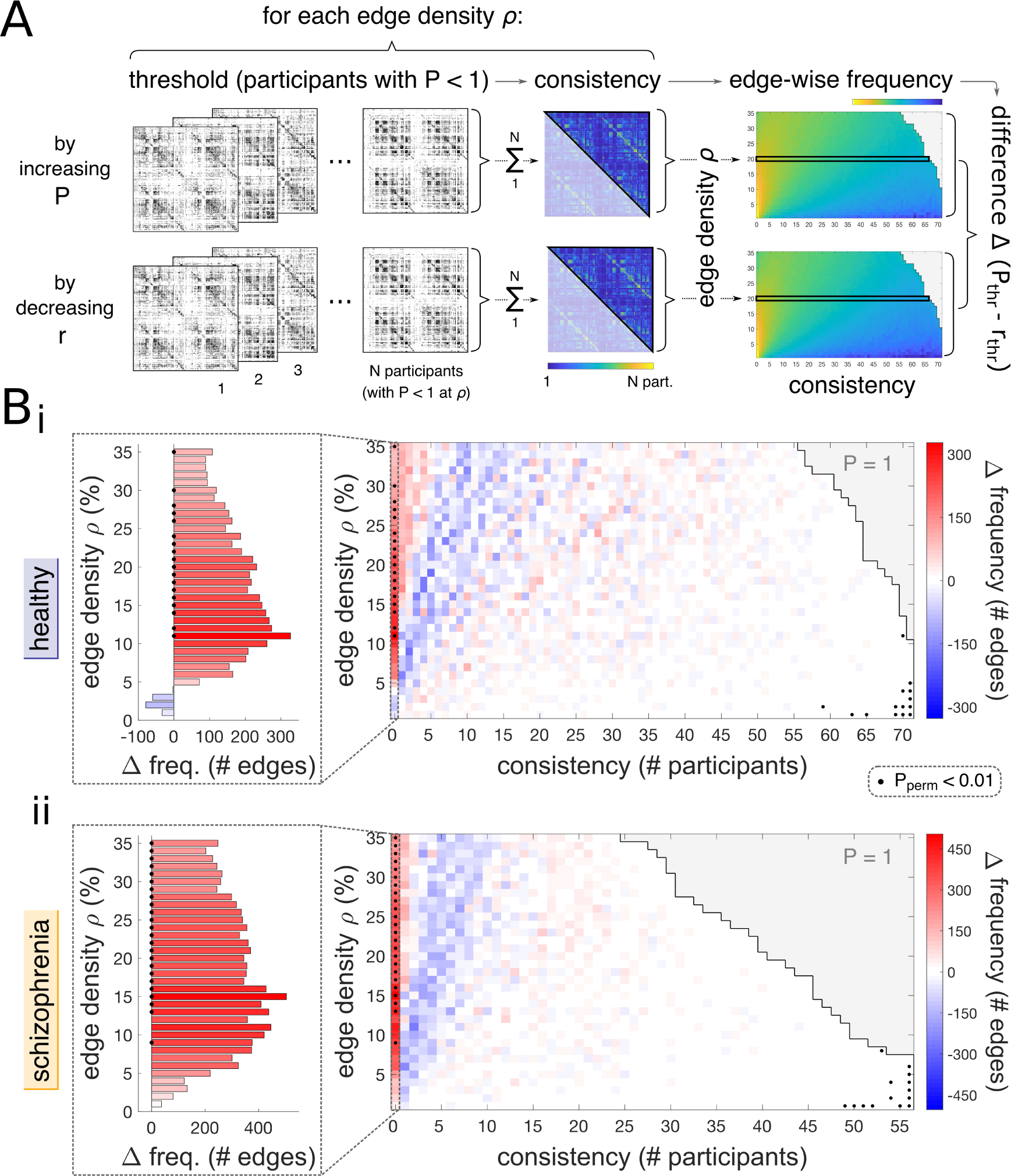
P-thresholded connectomes show increased edge consistency compared to r-thresholded connectomes. A) At each edge density and for each thresholding method (r and P), we quantified the consistency of each edge by counting the number of times each edge was present across the group at a given density. We then counted the number of edges that were found to occur at each level of consistency (i.e. the frequency of edges as a function of consistency). We did so for both r- and P-thresholding methods, and then evaluated the difference between the two methods. We estimated the significance of these differences by repeating the above procedure 10’000 times following permutation or r- and P-thresholded networks within group; P values were estimated as the proportions of empirical differences surpassing the permuted ones. As in previous analyses where networks were thresholded by fixed P value, we only considered participants with edge P < 1 at each density. B) Connectomes thresholded by decreasing P value showed an overall increased consistency of edges compared to connectomes thresholded by decreasing r. This was evidenced by an increased frequency of edges absent in all participants in P-thresholded networks (left inset histograms), in both (i) healthy controls and (ii) patients with schizophrenia.

#### 3.7 (In)dependence of results on motion

To confirm that the wavelet despiking algorithm had satisfactorily corrected the fMRI time series for effects of head micro movements during scanning, we demonstrated that correlations between pairs of regional fMRI time series showed no distance-dependent artefacts (Satterthwaite et al. (2012); SI Fig. S8A).

We also found that there was no significant difference between groups in the global mean effective *df* (r = - 0.16, P_MWU_ = 0.12; SI Fig. S8B). The median of the mean effective *df* across patients with schizophrenia was 36.4, [Q1,Q3] = [35.7,36.7], while the median of the mean effective *df* across healthy controls was 36.6, [Q1,Q3] = [36.2,36.7]. Moreover, there were no significant between group differences in nodal effective *df* (SI Fig. S8C; all FDR-adjusted nodal P_MWU_ > 0.12). These results indicate that, in both groups, movement-corrected fMRI time series have considerably lower effective *df* than their nominal *df*; but the absence of between-group differences in effective *df* indicates that the severity of head movement artefact, and therefore the number of *df* lost to wavelet despiking, was not significantly different between healthy controls and patients with schizophrenia.

Finally, although node degree is correlated to regional *df* within participants (median [Q1,Q3] Spearman *ρ* = 0.23 [0.11,0.34] for healthy controls, 0.21 [0.08,0.36] for patients with schizophrenia), as already reported for node strength in (Patel and Bullmore, 2016), there is no difference in the extent of these relationships between the two groups (rank-biserial *df* = -0.036, P = 0.73; Fig. S9C).

### 4. Discussion

In this article we demonstrate novel applications for probabilistically-thresholded functional connectomes based on edge-specific P values adjusted for loss of degrees of freedom (*df*) due to motion (Patel and Bullmore, 2016). We apply these methods to a population of healthy controls and patients with schizophrenia, and demonstrate reduced edge density and increased disconnectedness in patient connectomes when thresholding to variable density by P value (at P_FDR_<0.01). We confirm that comparison of functional connectomes with different weight distributions (such as patients with schizophrenia and controls) using arbitrary weight-based thresholds risks introduction of “non-significant” false positive edges and thus group differences in topological organisation (Fornito et al., 2012; van den Heuvel et al., 2017). The probabilistic thresholding methods we propose to mitigate this risk are statistically principled and simple to implement. We further demonstrate that only few individual connectomes can be constructed to the typically studied densities of 5-30%. Routinely thresholding to these densities without accounting for edge probabilities increases the risk of type I error. Using the wavelet-despiking-based probabilistic connectivity methods we describe additionally allows identification of the maximum connection density to which a connectome can be built to enable control over false positive results. Finally, we explicitly compare the application of thresholding connectomes by P value (following adjustment for effective *df*) to existing thresholding methods based on correlation (equivalent to assuming nominal *df*), demonstrating that (i) different edges are retained, (ii) these lead to differences in topology and (iii) the edges retained based on increasing P value are more consistent across subjects than edges retained based on decreasing correlation.

#### 4.1. Thresholding methods

Studies of functional connectome topology are typically conducted by retaining edges above a fixed weight (absolute thresholding), or by retaining a fixed proportion of the strongest edges (proportional or fixed-density thresholding; Fornito et al. (2013)). Of the two, fixed-density thresholding is most popular, likely because many measures of topological organisation depend on the edge density of the underlying graph (van Wijk et al., 2010). Furthermore, a recent study showed that differences in topological organisation between males and females, and between younger and older participants, were more stable following “proportional” (fixed-density) thresholding than “absolute” (fixed-amplitude) thresholding; in the latter case between-group differences alternated in direction (sign) across thresholds, and the evolution of differences in topological measures as a function of threshold magnitude was non-monotonic (Garrison et al., 2015).

However, it is known that thresholding to fixed edge density can lead to confounding effects when comparing groups with different distributions of edge weights – in a group with higher edge weights, this would lead to ignoring potentially important edges, while in a group with lower edge weights, this would lead to including weak or “nonsignificant” edges (van Wijk et al., 2010).

Fixed-amplitude thresholding is far less popular than fixed-density thresholding, having primarily been applied in earlier studies of functional connectivity (e.g.: Liu et al. (2008); van den Heuvel et al. (2009)). However, if applied in a principled manner, it can yield simple and interpretable graph-theoretical measures such as edge density, which might hold diagnostic promise. In a previous study of functional connectivity in schizophrenia, Alexander-Bloch et al. (2010) noted that the topological (dis)connectedness of a graph occurs at different (absolute) amplitudes of correlation for patients with schizophrenia and healthy controls, and remarked that this difference might have diagnostic potential. Accordingly, we find group differences within the edge density and disconnectedness of connectomes thresholded using fixed P values.

We note that there may have been previous studies of functional connectomes where association matrices were thresholded based on P values; however, these have assumed nominal *df* (number of points in the time-series), which we know to be an over-estimate of the effective *df* (Patel and Bullmore, 2016). Additionally, in a graph where all nodes possess equal *df*, the relationship between Pearson correlation r (the most common measure of association between regional neurophysiological time series) and the corresponding P value is perfectly monotonic (that is: as r increases, P decreases, and a given value of r maps to a single value of P). However, given the inhomogeneity of artefacts in space and time, we know that the relationship between r and P cannot be perfectly monotonic after denoising, because the loss of *df* is not spatially or temporally homogeneous (Patel and Bullmore, 2016). Wavelet despiking enables construction of probabilistic connectomes after quantifying the inhomogeneous loss of *df*. This is what gives rise to differences in topological organisation between functional connectomes thresholded (to fixed density) based on correlations or df-corrected P values. Although only small proportions of edges differ between connectomes thresholded by increasing *df*-adjusted P value or decreasing correlation (up to ~10%, depending on the participant and edge density), these differences are sufficient to lead to differences in topological organisation. The probabilistically-thresholded connectomes display reduced efficiency and generally increased transitivity (at higher densities), and thus reduced randomness, consistent with the exclusion of “noisier” edges at any given edge density. Further, the probabilistically-thresholded networks also show increased consistency across subjects.

Beyond hard thresholding to arbitrary densities, global probabilistic methods have been proposed which estimate dependence of the minimum absolute threshold on the number of samples (fMRI frames) and the number of nodes, whilst maintaining appropriate global FWER control (De Vico Fallani et al., 2014). Other methods include integrating across a range of arbitrary thresholds (Ginestet et al., 2011), or attempting to choose thresholds in an informed, non-arbitrary manner (De Vico Fallani et al., 2017) based on the cost-efficiency trade-off (Bullmore and Sporns, 2012). Methods presented herein extend beyond global probabilistic and principled thresholding methods by taking into account the spatially and temporally inhomogenous loss of *df* and thus the heterogeneity of correlation probabilities across the brain.

It is worth stating that thresholding is a difficult problem. While it is possible to estimate type I error (false positive rate) based on some assumptions on the statistical properties of the data, due to the lack of ground truth, it is difficult to assess type 2 error (false negative rate) in resting-state data. Thus, while it seems sensible to eliminate non-significant estimates of functional associations between regions, very tight control over type I error (and thus thresholding to very sparse densities) may theoretically lead to increased type 2 error and removal of signal. Indeed, weak correlations may contain diagnostic information, specifically with respect to schizophrenia (Bassett et al., 2012). Thus, “soft thresholding” methods designed to suppress rather than remove weaker connections (Schwarz and McGonigle, 2011), and methods to analyse unthresholded (fully connected) weighted connectomes (Rubinov and Sporns, 2011) have much value. Moving forward, wavelet-despiking based methods could be extended to unthresholded graphs by providing a *df*-based temporal weighting for each regional time series, thus effectively adjusting for loss of *df* and detection power upon estimation of association between regional neurophysiological time series.

#### 4.2. Topological randomisation and schizophrenia

A number of studies have reported increased topological randomisation of functional connectomes in patients with schizophrenia (Rubinov et al., 2009; Alexander-Bloch et al., 2010; Lynall et al., 2010; Lo et al., 2015). However, as suggested by Fornito et al. (2012), this was likely a consequence of applying fixed-density thresholds to groups with different edge weight distributions: “in the presence of a global reduction of mean functional connectivity in patients […] any analysis of graphs matched for connection density, *κ*, will result in the inclusion of proportionally more low-value (non-significant) edges in patients’ networks. If these values merely reflect noise, their inclusion will produce a more random topology.” Methodologies such as those we propose here, where cohorts are matched by connectivity probabilities, can be used to control for this.

When using traditional density thresholding, comparing cohorts regardless of differences in weight distributions, we found increased global efficiency in patients with schizophrenia relative to controls. With progressive randomisation, the path length of a graph decreases (Watts and Strogatz, 1998) while its global efficiency increases (Latora and Marchiori, 2001); this finding therefore aligns with previous reports of increased “randomisation” in schizophrenia. However, this effect disappears when controlling cohorts for type 1 error, i.e. when excluding connectomes which contain edges with probabilities P_FDR_ > 0.01 (and thus non-significant edges) from analysis. We further show, by comparing within each cohort connectomes with nonsignificant edges (P_FDR_ > 0.01) and those without, at a range of edge densities, that differences in global efficiency are driven by inclusion of non-significant edges. In other words, inclusion of noise increases efficiency and produces a more random topology. These findings remained qualitatively consistent when normalising with respect to random networks with preserved degree distributions (Rubinov and Sporns, 2010).

Together, these findings confirm the hypothesis advanced by Fornito et al. (2012) that increased randomisation in schizophrenia results from correlation-based thresholding to fixed edge density. At a lower level, these differences seem to be driven by a noise-driven “coalescence” of the fixed-density patient connectomes relative to healthy ones. When all participants are compared (regardless of edge significance), patients with schizophrenia exhibit lower numbers of (larger) connected components, in line with a previous study using fixed-density correlation-based thresholding (Bassett et al., 2012). However, these differences are substantially less extensive when only connectomes with edge P_FDR_ < 0.01 are included in analysis. Accordingly, connectomes with non-significant edges exhibit lower numbers of (larger) connected components. As the topological path length between pairs of disconnected nodes is infinite, the corresponding contribution to efficiency (which depends inversely on the path length) will be null, and the global efficiency will decrease (Fornito et al., 2016). The lower number of connected components in non-significant connectomes suggests that randomly placed non-significant edges are likely to act as “bridges” between disparate clusters of significant edges.

In line with previous reports of decreased segregation in functional connectomes of patients with schizophrenia, reported as decreased local efficiency (Liu et al., 2008) and clustering in both patients (Alexander-Bloch et al., 2010; Lynall et al., 2010) and their siblings (Lynall et al., 2010), we found decreased transitivity (a global measure of segregation) - although only at certain edge densities. Still, general decreases in transitivity in non-significant connectomes (both within healthy controls and patients with schizophrenia) suggest that the addition of noisy edges increases topological randomness by increasing transitivity. When normalising with respect to random networks with preserved degree distributions, we found increased normalised transitivity in patients relative to controls, and in connectomes with non-significant edges relative to those without non-significant edges. These disparities between our findings and the literature might be explained by the application of different measures – whereas the widely used average clustering coefficient has a dependence on the degree distribution of the underlying graph, transitivity does not (Newman, 2010). Furthermore, we can speculate that transitivity might be less affected by noisy edges than efficiency, as the addition of a single “noisy” edge will affect all paths traversing that edge, whereas it will only affect transitive closure in the immediate vicinity of the nodes it is connected to.

Our findings on topological randomisation align with a recent study by van den Heuvel et al. (2017), which demonstrates that group differences in overall functional connectivity strength leads to group differences in topological organisation of fixed-density connectomes, across several empirical and simulated case-control datasets. We note a relationship between our experiments and those in van den Heuvel et al. (2017): when comparing subsets of significant participants to ensure control over type I error, the difference in mean correlation between healthy controls and patients with schizophrenia is reduced. Subsequently, subsets of significant and non-significant participants show large differences in mean correlation. van den Heuvel et al. (2017) postulate possible strategies to control for effects of group differences in average functional connectivity, including the use of regression and permutation testing (van den Heuvel et al., 2017).

Here we describe a statistically-reasoned approach to limit inclusion of non-significant edges, which may result in artefactual group differences in topological organisation. This methodology, based on the probabilistic connectivity method described in Patel and Bullmore (2016), generates estimates of effective *df* from robust denoising of motion artefacts by wavelet despiking (Patel et al., 2014). To the best of our knowledge, wavelet-based methods used here (Patel et al., 2014; Patel and Bullmore, 2016) are the only currently available means of estimating the spatial and temporal variability in loss of confidence in time series recorded from motion-affected brain areas. As discussed in Patel and Bullmore (2016) and as demonstrated here on a dataset of healthy controls and patients with schizophrenia, this loss of effective *df* has substantial effects on the estimates of topological organisation of functional connectomes.

#### 4.3. Recommendations

Below, we offer guideline suggestions relating to graph construction and analysis, without focusing on denoising or other pre- or post-processing steps (although these should be carefully considered). Moreover, while we focus on Pearson’s correlation as a measure of association between regional time-series, the following guidelines also apply to other methods of connectivity estimation. Finally, while this paper focuses on functional brain networks, most of the following points are also relevant to structural brain networks, as further discussed below. We recommend the following with respect to network construction:

1. If studying functional networks, we advise careful consideration of time series frequencies analysed. Here we show that scale 4 frequencies (0.015-0.03 Hz) do not contain enough *df* after signal denoising to make any meaningful inference on associations between pairs of time series. Inferences at these low frequencies are likely to be erroneous.
2. Prior to the application of any potential threshold, the full weight distributions should be characterised. In particular, effects of mean connectivity, such as group differences or effects of age (depending on the question of interest of the study) should be well understood.
3. The application of methods for the characterisation of fully weighted, unthresholded networks (e.g. Rubinov and Sporns (2011)) should be considered.
4. If thresholding, we advise doing so according to some criterion of connection ‘likelihood’, rather than connection weights (e.g. correlations). The probabilistic method presented herein is an example of such a criterion, for the specific case of fMRI correlation networks. If studying functional networks which include negatively weighted edges, be aware of the potential effects on topological organisation of including these.
5. The choice of threshold should ideally be principled. Probabilistic thresholding lends itself well to a principled choice of threshold, based on statistical significance. In the present study, we retained edges with P_FDR_ < 0.01. Still, it is worth verifying that results are not exclusive to a single threshold.
6. In networks thresholded to variable edge density (e.g. by fixed P value, such as P_FDR_ < 0.01 used herein), “higher-order” graph theoretical measures (such as efficiency) will retain a dependence on edge density (van Wijk et al., 2010). In this case, it may be preferable to focus on simpler graph measures such as edge density itself, or the architecture of connected components.
7. For density-based thresholding, we would advise identifying what proportion of subjects contain significant or non-noisy edges at each edge density. At lower edge densities, a greater proportion of connectomes will contain only significant edges.
8. If wishing to enforce node-connectedness, the minimum-spanning tree method can potentially be applied (Alexander-Bloch et al., 2010); however, this risks the inclusion of a large proportion of false-positive edges, particularly at low edge densities. Therefore, the impact of such steps should be investigated.
9. It is worth investigating the relationship between any graph-theoretical measure used and mean connectivity. Previous work indicates that this might be weaker at lower edge densities (van den Heuvel et al., 2017). Investigation of this relationship will ensure that any effects reported in graph-theoretical measures are not driven by potential underlying effects of mean connectivity.

We note that many of the above suggestions are relevant to the thresholding of structural networks as well. For example, the fact that inclusion of noisy edges may lead to erroneous conclusions is pertinent to the case of structural connectivity derived from diffusion imaging (Zalesky et al., 2016). Moreover, various attempts have been made in the spirit of these recommendations to retain connections based on some criterion of connection “likelihood” rather than simply assuming that the strongest edges are the most likely. Examples include identification of edges which are consistently detected (de Reus and van den Heuvel, 2013) or consistently strong (Roberts et al., 2017) across participants in structural networks derived from diffusion imaging, or edges which are consistently strong across bootstrapped samples of participants in structural correlation networks (Váša et al., 2017).

Following the above recommendations should help safeguard against the introduction of potentially artefactual group differences in topological organisation in future analyses of brain connectivity.

#### 4.4. Further considerations and future work

We note that the methods of adjusting regional *df* and consequently edge-wise P values for the effects of spatially (and temporally) heterogeneous denoising using wavelet despiking, proposed in Patel and Bullmore (2016) and applied here, do not provide a universal solution to dealing with all noisy data. The *df* correction is needed to remove bias in interpretability of connectivity values; that is, to disentangle whether low connectivity is due to low *df* or due to low intrinsic connectivity (as discussed in Patel and Bullmore (2016)) and subsequently to control for type I error. While application of wavelet despiking will denoise regional time-series and subsequent adjustment of P values for effective *df* will take the effects of denoising into account when thresholding, these methods (like all other methods) are unable to “restore” the lost signal in these regions. Thus, users of the methods presented herein should remain aware of potential biases, such as (i) group differences in average or regional *df* (not present in the current analyses), (ii) within-subject relationships between regional df and connectivity (present in the current analyses, but not different between groups) or (iii) exclusion of a different number of subjects from each group (whether due to high motion at the beginning of the analysis pipeline, or low edge probabililty when thresholding to high edge densities), leading to potential imbalances in sample size between groups. Taken together, the methods applied herein enable mitigation of the effects of motion on functional connectomes; however, they do not offer a panacea to attempted analysis of low quality data – whether due to high motion, low SNR or other reasons.

The ability to robustly estimate associations between the activity of pairs of regions depends on factors other than motion. Perhaps the most important of these is the length of the time series. Earlier reports suggested that estimates of functional correlation strength stabilise within five minutes (Van Dijk et al., 2010) and that graph-theoretical measures of functional connectome organisation stabilise only after two minutes (Whitlow et al., 2011). However, more recent studies suggest that reliability does substantially increase with increasing scan duration, plateauing at 12 minutes (Birn et al., 2013) with increased reliability of 14 minutes relative to 7 minutes (Termenon et al., 2016). A recent meta-analysis confirmed increasing test-retest reliability of graph-theoretic fMRI studies with increasing scan duration, especially beyond five minutes (Andellini et al., 2015). Interestingly, whole brain reliability seemed stable with increasing sampling rate for scans of equal duration (Liao et al., 2013), suggesting that the duration of the acquisition has importance above and beyond the sampling rate. From a purely statistical point of view, the larger the number of samples (either longer time series or faster sampling rate), the larger the number of effective *df*; this should translate to a higher density a graph can be built to while ensuring control over type 1 error.

We also note that the dependence of network topology on factors such as scan duration or (in the case of P value thresholding) the significance level (e.g.: *α* = 0.05, 0.01 or 0.001) precludes us from claiming that there exists a single “true” functional network topology that our method is able to recover – rather, the topology will be conditional on such factors. Still, we believe that in limiting the inclusion of spurious edges, our method comes closer than existing thresholding approaches to estimating the accurately denoised topology. Similarly, we do not claim that there are definitely no differences in topological organisation between patients with schizophrenia and healthy controls, even in graphs thresholded to fixed edge density by *df*-adjusted P value, simply on the basis of our rejection of the null hypothesis. However, it seems likely, based on our results and those of van den Heuvel et al. (2017), that the previously reported differences were methodologically grounded in the thresholding of networks to fixed edge density based on correlation – and we suggest that this can be avoided by using the methods presented in this paper.

Additionally, loss of power might arise from the averaging of voxel-wise signals over functionally inhomogeneous regions of interest. Random sub-parcellations of anatomical atlases (Cammoun et al., 2012; Romero-Garcia et al., 2012), as used here, lead to hundreds of regions whose small size should limit the spatial blurring of functional boundaries. Data-driven parcellations might improve the ability to detect changes in functional connectivity across conditions and development (Yeo et al., 2011; Glasser et al., 2016).

Furthermore, new multi-variate methods of inter-regional association are being developed which appear more accurate than simple correlations (Geerligs et al., 2016); extending such methods to take into account the regionally heterogeneous loss of degrees of freedom due to motion and other scanning artifacts (Patel and Bullmore, 2016) should yield ever-more precise estimates of functional connectivity.

Analysis of dynamic variability of functional connectivity is an increasingly popular avenue (e.g.: Zalesky et al. (2014); Karahanoglu and Van De Ville (2015)) that we did not explore here. In this context, the importance of denoising methods that preserve the structure of neurophysiological time-series (Kundu et al., 2012; Patel et al., 2014), and can further adjust estimates of the optimal sliding-window size (Leonardi and Van De Ville, 2015; Zalesky and Breakspear, 2015) to ensure fixed effective *df* across time (Patel and Bullmore, 2016) should become increasingly clear. Methods such as frame censoring (scrubbing) (Power et al., 2012), widely used in the neuroimaging community, are unsuited to denoise artefacts in such cases as they disrupt the temporal structure of the BOLD signal.

Beyond altered connectivity, intrinsic properties of regional neurophysiological signal are altered in schizophrenia. Such alterations are found in the power (Zalesky et al., 2012; Yang et al., 2014) and entropy (Yang et al., 2015) of the BOLD signal. Interestingly, these alterations correlate to measures of functional dysconnectivity in patients with schizophrenia, although regional and inter-regional signal properties may be differentially sensitive to disease deficits (Bassett et al., 2012; Zalesky et al., 2012). Moreover, it is known, from studies using MEG, that inter-frequency coupling is affected in schizophrenia (Siebenhühner et al., 2013; Brookes et al., 2016); while fMRI bandwidth is too narrow for investigations of inter-frequency effects, recording methods with high temporal resolution such as EEG and MEG could equally benefit from processing and analysis using the methods discussed here.

Finally, the lack of clinical or behavioural measures prevented us from evaluating the true diagnostic relevance of the probabilistic thresholding methods presented here. Studying such relationships is a necessary target of future work, if fMRI connectomics are to become clinically useful (Matthews and Hampshire, 2016). An important question is whether probabilistically thresholded functional connectomes lead to stronger relationships between measures of graph (dys)connectivity and cognitive, behavioural or clinical scores. Still, given the likely multivariate nature of relationships between brain connectivity and cognition (Mišić and Sporns, 2016), identifying such relationships is non-trivial.

#### 4.5. Conclusions

In summary, probabilistic thresholding of denoised functional connectomes at fixed P value confirms greater dysconnectivity in schizophrenia, with fewer significant connections between regions. Thresholding to fixed edge density (by effective-df-adjusted P value) supports the view that previously reported increases in randomisation within functional connectomes of patients with schizophrenia were linked to inclusion of noisy edges within the patients’ connectomes, which exhibit shifted correlation distributions. Finally, while only a small proportion of edges differ between functional connectomes thresholded to fixed edge density based on edge weights and P values, these edges may lead to significant alterations in global topological organisation. Our results warrant care during analysis of connectomes constructed using correlations, and emphasise the need for exploration of potential “lower-order” causes (such as shifts in edge weight distributions between groups or across participants) of alterations in “higher-order” topological properties of brain network organisation. The code to denoise fMRI data using wavelet despiking, extract (regionally heterogeneous) effective df and adjust edge-wise P values for these is available at www.brainwavelet.org (BrainWavelet Toolbox v2.0), which will enable readers to implement the methods presented in this article in their analyses.

## Author contributions

Conceived and designed analyses: FV, ETB, AXP. Processed and quality controlled data: AXP. Conducted analyses: FV. Wrote the manuscript: FV, AXP. Critically appraised the manuscript: FV, ETB, AXP.

## Acknowledgments

We would like to thank Sarah Morgan, Alex Fornito and Andrew Zalesky for helpful discussions. FV was supported by the Gates Cambridge Trust.

## Competing interests

ETB is employed half-time by the University of Cambridge and half-time by GlaxoSmithKline; he holds stock in GSK.

## Availability of data and code

Raw anatomical and functional scans were kindly made available by the Mind Research Network and University of New Mexico (http://fcon_1000.projects.nitrc.org/indi/retro/cobre.html). Code to perform wavelet despiking, estimation of effective *df* and subsequent adjustment of P values is available in the Brain Wavelet toolbox (v2.0), at www.brainwavelet.org. Processed data, including correlation and *df*-adjusted P value matrices for all participants, has been uploaded to the Cambridge Data Repository (http://doi.org/10.17863/CAM.12827). Matlab code used to conduct main analyses is available from FV’s github: https://github.com/frantisekvasa/probabilistic_connectome (DOI: 10.5281/zenodo.847604).

